# Decoy Receptor Fine-tunes Cytokinin Signaling

**DOI:** 10.1101/2022.10.20.513092

**Authors:** Michaela Králová, Ivona Kubalová, Jakub Hajný, Karolina Kubiasová, Michelle Gallei, Hana Semerádová, Ondřej Novák, Eva Benková, Yoshihisa Ikeda, David Zalabák

**Author notes:** These authors contributed equally to this work (Michaela Králová, Ivona Kubalová).

## Abstract

Hormone perception and signaling pathways play a fundamental regulatory function in cell growth, developmental, and physiological processes in both plant and animal systems. Those pathways are activated by hormone binding to the receptor to trigger cellular responses. Equally important are mechanisms that suppress activated transduction cascades to reset the system. Different mechanisms at the level of hormone biosynthesis and deactivation through degradation, conjugation, and production of repressors that attenuate transduction cascades downstream of receptors are known. In animal systems, decoy receptors have been identified as another important mechanism for fine-tuning the activity of the signaling pathways in processes like inflammatory responses, apoptosis, and blood vessel formation. Decoy receptors recognize and bind specific signaling molecules, but they cannot activate downstream signaling pathways thus providing competitive inhibition. Here we describe the first decoy receptor in plants. We show that the splicing variant of *CRE1/AHK4* receptor of cytokinin, a hormone with a key role in the regulation of cell division and meristem maintenance in plants, acts as a decoy receptor to attenuate cytokinin signaling. We propose that this novel mechanism of signaling control applies in processes when modulation of CK signaling is needed.

## Introduction

Cell signaling is a fundamental feature of all living organisms and allows them to respond promptly to extracellular or intracellular signals. The signal inputs can be of either chemical nature ligands (hormones or other small molecules) or physical (e.g., temperature, light, pressure). A signal is perceived by receptors and transduced via the activated signaling cascade to trigger cellular responses (Kholodenko, 2006; Takeuchi et al., 2021).

The signaling pathways are stringently controlled at several levels via negative feedback loops as a part of the primary responses. The role of negative feedback loops is to reset the signaling and thus allow for prompt, strong, and temporary responses. A typical component of negative feedback regulation is the expression of a repressor attenuating the signaling pathway. At the receptor level, ligand accessibility to the receptor is controlled by its biosynthesis and degradation allowing for turning the signaling on and off (Perrimon and McMahon, 1999; Gordon et al., 2009; Ferrell, 2013; Lemmon et al., 2016).

Decoy receptors (DcRs) represent a unique negative feedback mechanism that evolved in mammalian systems (Mantovani et al., 2001). DcRs are capable of ligand binding but structurally unable to activate the signaling cascade. The DcRs compete with canonical receptors for ligand binding and thus attenuate the signaling cascade. A typical representative of DcRs is Interleukin 1 receptor type II (IL1R2) counteracting the IL1R1 receptor to attenuate the inflammatory reaction by competing for interleukin binding (Re et al., 1994). Decoy Receptor 3 (DcR3) titrates the TNF ligand (tumor necrosis factor) and thus prevents its binding to the canonical TNF receptors to modulate apoptosis (Ashkenazi, 2002). The activity of VEGFR1 (Vascular endothelial growth factor receptor 1), another DcRs is essential for angiogenesis and vascular growth (Meyer et al., 2006). In plants, decoy receptors, as an alternative mechanism to control the activity of hormonal pathways, have not been identified so far.

Cytokinins are key plant hormones and regulators of cell proliferation, meristem activity, shoot, and root branching, root vascular development, and chloroplast maturation, as well as of response to biotic and abiotic stresses (Miller et al., 1955; Werner and Schmülling, 2009).

The *Arabidopsis thaliana* genome encodes for three cytokinin receptors: *ARABIDOPSIS HISTIDINE KINASE 2* (AHK2), AHK3, and *AHK4/CYTOKININ RESPONSE 1 (CRE1)/WOODEN LEG (WOL*) (Inoue et al., 2001). The cytokinin signaling cascade is initiated upon ligand binding to the N-terminal CHASE (<u>c</u>yclases/<u>h</u>istidine-kinase-<u>a</u>ssociated <u>s</u>ensor <u>e</u>xtracellular) domain of the receptor leading to a conformational change allowing for ATP binding in the histidine kinase domain. The activated histidine kinase domain transfers a phosphate to the conserved aspartate residue within the C-terminal receiver domain. Five AHP (ARABIDOPSIS PHOSPHOTRANSMITTER1 to 5) proteins then transmit the phosphate from the receiver domain of the receptor to one of eleven type-B RESPONSE REGULATORS (B-ARRs) (Sakai et al., 1998; Sakai et al., 2000; Lohrmann et al., 2001; Hosoda et al., 2002). Phosphorylated B-ARRs trigger the expression of primary response genes.

The cytokinin signaling pathway is tightly controlled through several negative feedback loops. Type-A ARRs (A-ARRs) are induced by cytokinin (Brandstatter and Kieber, 1998; Sakakibara et al., 1998), and although they can receive a phosphate from activated AHPs they cannot trigger gene expression thus providing a competitive inhibition of B-ARRs (To et al., 2004). Other negative feedback mechanisms involve protein stabilization of A-ARRs upon cytokinin treatment (To et al., 2007), S-nitrosylation of AHP1 blocking its activation (Feng et al., 2013), proteasome-dependent degradation of type-B ARR (ARR2) by KMD (KISS ME DEADLY) ubiquitin E3 ligase complex (Kim et al., 2012; Kim et al., 2013), and induction of cytokinin degrading enzyme CKXs (CYTOKININ OXIDASES) (Werner et al., 2003; Werner et al., 2006). AHP6 uses its pseudo phosphotransfer domain to stop the phosphorelay (Mähönen et al., 2006b). All these mechanisms provide several layers of regulation to tightly control cytokinin signaling and allow for normal plant organ growth and development.

While *CRE1/AHK4* plays an essential role in root vascular development (Mähönen et al., 2000), the other two known cytokinin receptors are expressed mostly in the rosette leaves and inflorescences and to a lesser extent in the root tissue (Higuchi et al., 2004; Mähönen et al., 2006a). All three cytokinin receptors play a partially redundant function *in planta* (Riefler et al., 2006).

The cytokinin receptors are localized to the plasma membrane or endoplasmic reticulum (ER) (Kubiasová et al., 2020) with a ligand-binding domain facing the apoplast or ER lumen and a histidine kinase domain and a receiver domain exposed to the cytosol (Lomin et al., 2018). Point mutations in histidine kinase and receiver domains interfere with protein activity proving they are critical for receptor function (Mähönen et al., 2006a).

The most common AS event *in planta* is intron retention (IR) (Filichkin et al., 2010) often induced by abiotic stresses (Reddy et al., 2013). Nevertheless, the biological meaning of IR is still elusive. In mammalian systems, most intron-retaining transcripts are recognized by the nonsense-mediated mRNA decay pathway (NMD pathway) and targeted for degradation since their detrimental function represents a potential risk for the organism (de Lima Morais and Harrison, 2010). In plants, several IR transcripts escape the NMD-mediated degradation pathway, suggesting they might have some biological functions (Kalyna et al., 2012). One of the most important questions is whether the pool of IR transcripts is translated into protein or represents transcripts “on pause”, waiting for translation in the nucleus, once the intron is spliced out. So far only a few studies are showing that IR generates novel protein variants to fine-tune plant hormone signaling pathways (Wang et al., 2015; Zhan et al., 2015).

We propose a novel regulatory feedback mechanism of cytokinin signaling mediated by IR of the seventh *CRE1* intron creating a decoy receptor. Having functional reporter lines expressing both fluorescently tagged canonical CRE1(full-length) and CRE1^int7^ (truncated) proteins in our hands, we investigated a novel regulatory feedback mechanism of cytokinin signaling mediated by alternative splicing of *CRE1* seventh intron. In addition, we observed subcellular localization of both CRE1 protein variants in their native expression domain.

## Results

When cloning a *CRE1* open reading frame we isolated novel transcript variant *CRE1^int7^* produced via alternative splicing of the seventh *CRE1* intron. The *CRE1* gene consists of ten exons, interrupted by nine intron sequences (Fig. 1A). The first intron of *CRE1* undergoes complex alternative splicing generating six splice variants annotated in the TAIR database (www.arabidopsis.org). While first intron splicing controls the CRE1 protein N-terminus, the retention of the seventh intron introduces a premature termination codon (PTC) resulting in a truncated protein lacking the receiver domain affecting the receptor function (Fig. 1A). The function of *CRE1^int7^* was tested using a wide range of biochemical, molecular biology, and cell biological approaches and compared with the canonical CRE1 receptor.

**Fig. 1.**
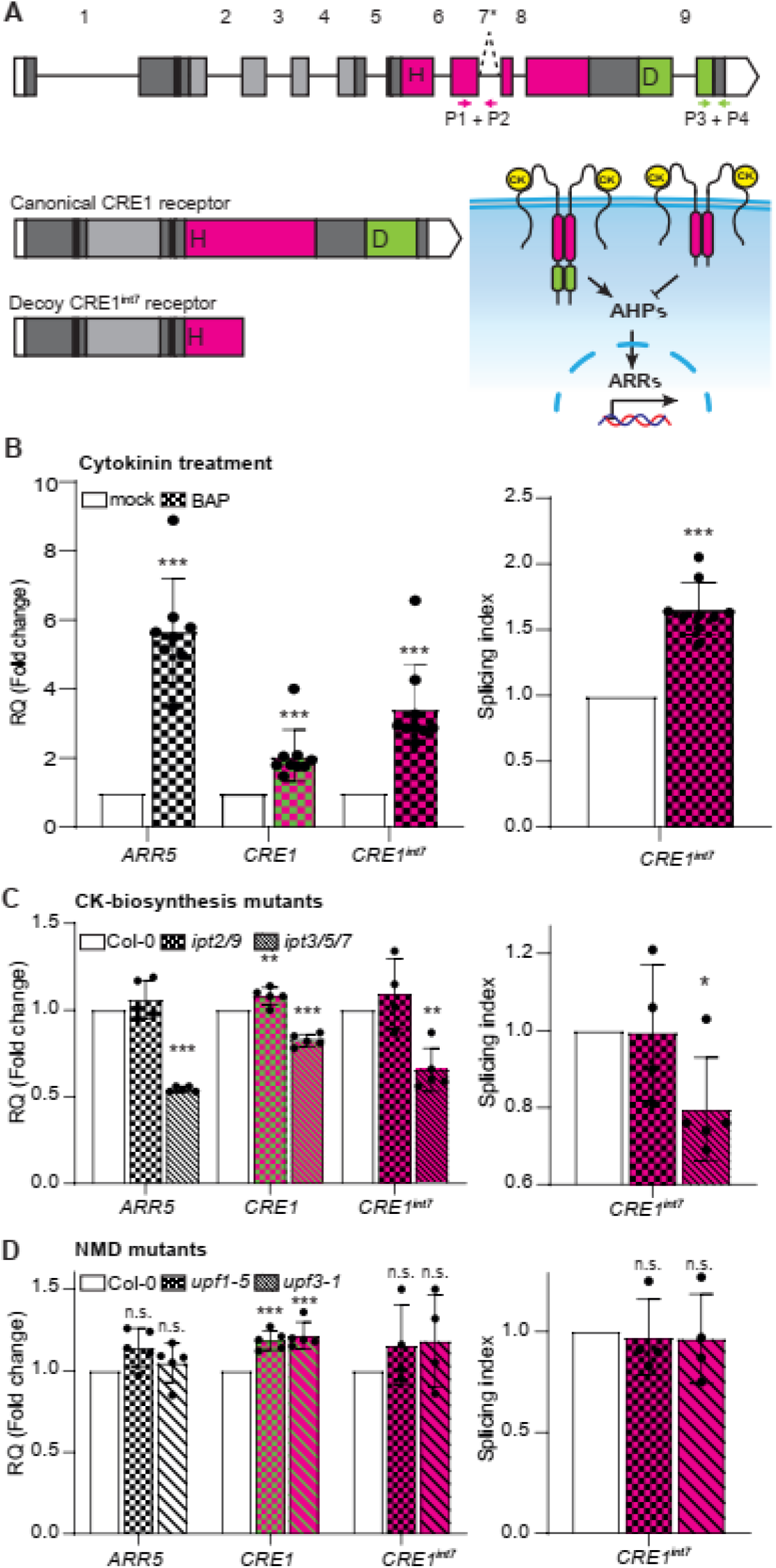
Alternative splicing of *CRE1* gene produces novel *CRE1^int7^* transcript; *CRE1^int7^* is responsive to cytokinin and is not a target to the NMD pathway. **A,** The *CRE1* gene consists of 10 exons and nine introns. The transcripts encoding the canonical CRE1 and decoy CRE1^int7^ receptors are produced through processes of constitutive or alternative splicing (intron retention). Retention (dashed line) of the seventh intron introduces premature termination codon (asterisk) into *CRE1* transcript, thus encoding the truncated receptors. The lack of the protein C-terminal part, including the conserved receiver domain, alters receptor function. The proposed modus operandi of canonical CRE1 and decoy CRE1^int7^ receptors is shown in the scheme (right). Upon cytokinin (CK) binding the canonical CRE1 phosphorylates AHPs and activates the signaling cascade. The phosphate is transmitted from AHPs to ARRs, ultimately triggering the expression of cytokinin primary responsive genes in the nucleus. The CRE1^int7^ lacking receiver domain cannot phosphorylate AHPs and thus act as a decoy to attenuate the signaling upon CK binding. *CRE1* transcripts encode ligand-binding CHASE domain (light grey), flanked by two transmembrane domains (black bars), histidine kinase domain (pink) with conserved histidine motif (H), receiver domain (green) with conserved aspartic acid (D). The 5’ and 3’-UTRs are shown in white. Primer pair position to detect *CRE1^int7^* (P1+P2) and overall *CRE1* transcripts (P3 + P4) used in RT-qPCR experiments (**B**). **B**, Quantitative RT-PCR of seventh *CRE1* intron retention upon CK (1 μM BAP, 1 h) application in Col-0 wild-type roots. **C**, Quantitative RT-PCR of seventh *CRE1* intron retention in roots of CK-biosynthesis mutants *ipt2/9* and *ipt3/5/7*. **D,** Quantitative RT-PCR of seventh *CRE1* intron retention in roots of nonsense-mediated RNA decay (NMD) mutants *upf1-5* and *upf3-1*. The relative gene expression (RQ) was normalized to *TUBULIN3* (left) in mock (**B**) or Col-0 wild-type roots (**C**, **D**). The splicing index of the seventh *CRE1* intron was normalized to overall *CRE1* transcripts (right). Values show the mean from nine (**B**) and five (**C**, **D**) independent experiments, respectively. The *P*-value shows the Student’s t-test, data comparing differences between mock and BAP or mutants (**B**) and Col-0 wild type (**C**, **D**). \**P* < 0.05, \*\**P* < 0.01, \*\*\**P* < 0.001, n.s., non-significant. Results represent the mean value ± standard error.

### Cytokinin interferes with splicing of the seventh intron

The *CRE1^int7^* transcript was isolated from Col-0 root explants cultivated on a shoot-inducing medium (SIM). Since the SIM contains a high cytokinin concentration, we hypothesized that the retention of the seventh intron within the *CRE1* transcript might be induced by cytokinin. To answer this question, we followed the expression of overall *CRE1* transcripts, and particularly of the seventh *CRE1* intron-retaining transcripts in *Arabidopsis* roots, using quantitative RT-qPCR in response to cytokinin 6-Benzylaminopurine, BAP (1 μM, 1 h). Primer pair annealing within the last exon was used to detect all *CRE1* transcript variants while pair annealing to the seventh exon and intron specifically recognizes the *CRE1^int7^* variant (Fig. 1A). The specificity of RT-qPCR amplicons was verified by sequencing. To exclude the possibility of contamination with genomic DNA, the isolated RNA was tested using primers binding to the non-transcribing promoter region of *ERF8 (Ethylene Response Factor 8*). To figure out the *CRE1^int7^* proportion within the *CRE1* transcripts pool we have evaluated the so-called splicing index. The splicing index is calculated as the relative expression of *CRE1^int7^* normalized to overall *CRE1* transcripts. Overall *CRE1* transcripts were significantly accumulated in roots one hour after cytokinin application compared to mock (DMSO, solvent) treatment (Fig. 1C). The *CRE1^int7^* transcripts accumulated even stronger in response to cytokinin. Indeed, the *CRE1^int7^* splicing index was significantly increased in CK-treated roots compared to mock (Fig. 1C), implying cytokinin interferes with the splicing machinery and impairs the splicing of the seventh *CRE1* intron leading to the accumulation of *CRE1^int7^* transcripts. *ARR5*, the cytokinin primary response gene, was used as a control for a cytokinin treatment.

To assess the effect of endogenous cytokinins on *CRE1^int7^* splicing, we employed the cytokinin biosynthesis mutants *ipt3/5/7* and *ipt2/9*. The levels of isopentenyl adenine and *trans*-zeatin (*t*Z), the most bioactive cytokinin forms, and their corresponding metabolites are decreased in *ipt3/5/7*. On the other hand, the mutant *ipt2/9* lacks tRNA-derived cytokinins, a less active *cis*-zeatin *c*Z-type (Miyawaki et al., 2006). Reduced cytokinin biosynthesis in *ipt3/5/7* led to dampened cytokinin activity, as demonstrated by a drop in *ARR5* expression compared to Col-0 wild-type roots (Fig. 1D). Similarly, the overall *CRE1* transcripts, as well as *CRE1^int7^*, were reduced in *ipt3/5/7* mutant roots compared to the Col-0 wild type. Moreover, the *CRE1^int7^* splicing index significantly decreased in the *ipt3/5/7* mutant (Fig. 1D). However, the *CRE1* expression and *CRE1^int7^* splicing index did not change in the *ipt2/9* double mutant. These results suggest that *CRE1^int7^* splicing is controlled by the level of active cytokinins.

To find out if *CRE1^int7^* transcripts have biological relevance, we have analyzed the expression level as well as the splicing index of *CRE1^int7^* in *upf1-5* and *upf3-1* loss-of-function mutants deficient in key components of the NMD machinery in *Arabidopsis* (Arciga-Reyes et al., 2006). Although *CRE1* expression is slightly increased in *upf1-5* and *upf3-1* mutants, the *CRE1^int7^* splicing index is unchanged (Fig. 1B), suggesting it may have a biological function.

### CRE1^int7^ can bind cytokinin but not activate the signaling cascade

To explore the consequences of seventh intron retention on the CRE1 receptor function we have analyzed the ligand-binding capacity of both CRE1 variants in the *E. coli* expression system (Romanov et al., 2005). This assay is based on competition of tritiated ligand [^3^H]*t*Z, specifically bound to the receptor protein, with unlabeled cytokinin ligands (iP, tZ). If the protein is capable of ligand binding, the [^3^H]*t*Z is displaced by cold iP and tZ, resulting in a CPU signal decrease. The CRE1^int7^ protein specifically bound cytokinin ligands [^3^H]*t*Z to a similar extent as the canonical CRE1 receptor. On the other hand, the control culture, carrying an empty expression plasmid, showed only a weak background signal, verifying the specificity of CRE1 ligand binding (Fig 2A). *t*Z and iP displaced comparably the [^3^H]*t*Z from both receptor proteins accompanied by decreased CPU levels (Fig. 2A). Next, we tested the receptor activation of both receptor variants upon cytokinin binding in *E. coli* (Suzuki et al., 2001). The canonical CRE1 strongly activated the expression of β-galactosidase reporter upon iP treatment, while CRE1^int7^ did not similar to the control culture carrying the empty vector (Fig. 2B). The expression of CRE1 receptor proteins in *E. coli* was verified by western blotting (Sup. Fig. 1A). These experiments showed the CRE1^int7^ variant retains the ligand-binding capacity, but it cannot transmit the signal downstream of the signaling cascade, due to the missing receiver domain.

**Fig. 2.**
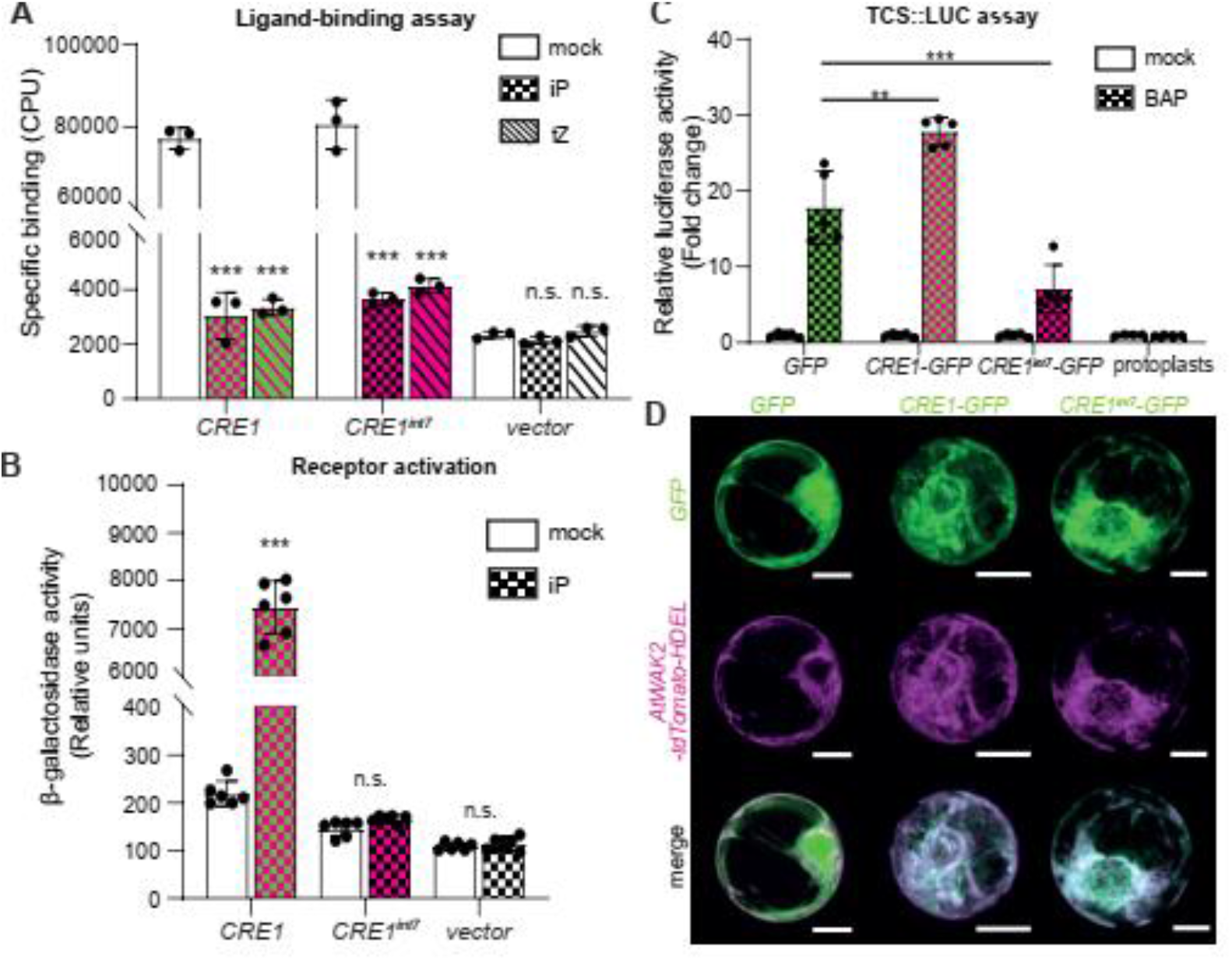
CRE1^int7^ binds cytokinin but cannot activate the signaling pathway. **A,** Competitive ligand binding assay in *Escherichia coli* expressing CRE1, CRE1^int7^, and empty vector *pIN-III* (vector). The binding of 3 nM [2-^3^H]*t*Z was assayed together with unlabeled 10 μM iP and tZ. Values show the mean from three biological replicates ± standard deviation. The *P*-value shows the Student’s t-test, data comparing differences between mock and iP or *t*Z. \*\*\**P* < 0.001, n.s., non-significant. **B**, Comparison of cytokinin response in *E. coli* KMI001 receptor activation assay with recombinant CRE1 and CRE1^int7^ receptors, triggered by 1 μM iP. The mock treatment represents solvent control DMSO (0.1%). The values represent the mean from six biological replicates ± standard deviation. The *P*-value shows the Student’s t-test, data comparing differences between mock and iP. \*\*\**P* < 0.001, n.s., non-significant. **C**, *TCS::LUCIFERASE (TCS::LUC*) cytokinin reporter activity in *Arabidopsis* Col-0 protoplasts expressing CRE1-GFP is significantly upregulated in response to cytokinin (0.5 μM BA) when compared to protoplasts expressing GFP alone. On the contrary, the *TCS::LUC* activity was significantly lower in protoplasts expressing CRE1^int7^-GFP. The values represent the mean from five biological replicates ± standard deviation. The Student’s t-test, data comparing differences between protoplasts expressing CRE1-GFP or CRE1^int7^-GFP, and GFP alone (*35S::GFP). **P* < 0.01, *** *P* < 0.001. **D**, Confocal microscopy of transiently expressed *35S::CRE1-GFP, 35S::CRE1^int7^-GFP*, and *35S::GFP* control in *Arabidopsis* protoplasts (green). Protoplasts were co-transformed with *35S::AtWAK2-tdTomato-HDEL* (ER marker, magenta). Maximum intensity projection of at least five Z-sections of protoplast is shown. Scale bars, 10 μm. At least 6 individual protoplasts were imaged.

### CRE1^int7^ attenuates cytokinin signaling by functioning as a decoy receptor

We have further tested the function of CRE1^int7^ using a *TCS::LUC* assay in *Arabidopsis* protoplasts, plant cells with removed cell walls (Müller and Sheen, 2008). The *TCS::LUCIFERASE (TCS::LUC*) is an established transient reporter system to monitor cytokinin output *in planta*. It consists of a synthetic promoter (*TCS*, two-component system), responsive to cytokinin, driving the expression of the firefly luciferase reporter gene (*LUC*). This system allows us to test the effect of studied proteins on the cytokinin signaling machinery in a plant cell context. The cytokinin readout is measured as LUC enzyme activity.

The CRE1 variants were C-terminally fused with *GFP* and expressed in protoplasts isolated from Col-0 root suspension culture. In the control sample, expressing only free GFP, the relative luciferase activity increased upon treatment (500 nM BAP/12 h), (Fig. 2C). This background activity is attributed to the native cytokinin signaling machinery present in protoplasts. The cytokinin response was further increased in protoplasts overexpressing the canonical CRE1-GFP receptor fusion. On the contrary, the overexpression of CRE1^int7^-GFP fusion protein diminished the relative luciferase activity in Col-0 protoplasts compared to the GFP control (Fig. 2C). This significantly weaker cytokinin response indicates that CRE1^int7^ can interfere with the native cytokinin signaling machinery present in Col-0 protoplasts to attenuate it. Both CRE1-GFP proteins co-localized well with the ER marker AtWAK2-tdTomato-HDEL in Col-0 protoplasts, (Fig. 2D). The localization pattern, as well as the signal intensity, was comparable between the CRE1-GFP and CRE1^int7^-GFP fusion proteins, implying both variants localize and function in the same compartment/s. The expression of the CRE1 proteins was further verified using Western blotting (Sup. Fig. 1B). Altogether, the results support the CRE1^int7^ function as a decoy receptor attenuating the CK signaling machinery.

### CRE1^int7^ attenuates cytokinin signaling in planta

To explore the function of both CRE1 variants *in planta* we generated transgenic *Arabidopsis* lines expressing the CRE1-GFP fusion proteins under the control of the 2 kbp native *CRE1* promoter (Michniewicz et al., 2015)(*pWOL*;. The construct encoding the canonical receptor CRE1-GFP was introduced into the *cre1-2* mutant background. Since CRE1^int7^ protein cannot activate the cytokinin signaling in bacteria (Fig. 2B), we expected CRE1^int7^-GFP would not complement the *cre1-2* mutant phenotype. Therefore, CRE1^int7^-GFP was brought into the Col-0 background. CRE1-GFP, as well as CRE1^int7^-GFP proteins, were strongly expressed in the root stele (Fig. 3A). This agrees with the previously published CRE1 expression pattern (Mähönen et al., 2006a). Both proteins are predominantly present in all tissues of root stele, including phloem cells, xylem cells as well as intervening cambial cells. The strong expression could be detected also during the early stages of lateral root formation, and it persists during lateral root emergence (Fig. 3B). The CRE1-GFP as well as CRE1^int7^-GFP fusion proteins showed ER localization pattern and accumulated at the PM in cells of the root stele. Moreover, the PM signal is polarized concentrating in the apical and basal sides of the root stellar cells (Fig. 3A).

**Fig. 3.**
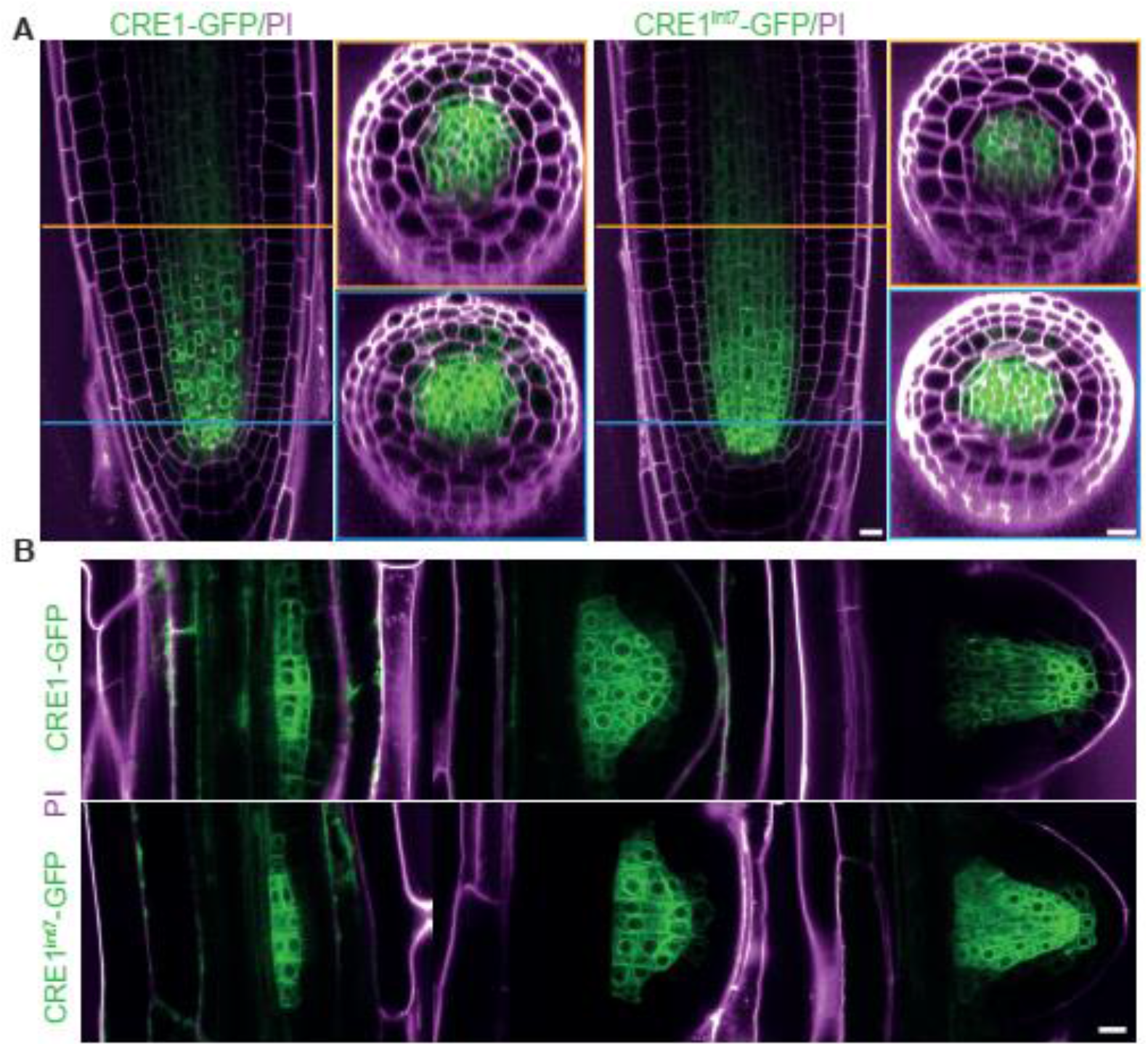
CRE1 and CRE1^int7^ are expressed in the root stele and lateral root primordia. **A,** Confocal microscopy of CRE1-GFP (left panel) and CRE1^int7^-GFP (right panel) 5-days-old seedlings. Longitudinal section through the root central zone (left), cross-sections through the root apical meristem in two positions (blue and orange square). Orange and blue horizontal lines show the position of the focal plane of individual root cross-sections. **B**, Confocal microscopy of CRE1-GFP (top panel) and CRE1^int7^-GFP (bottom panel) 6-days-old seedlings in primordia of three stages of developing lateral roots. Propidium iodide (magenta), GFP (green). Scale bars, 10 μm. At least 15 individual roots were imaged.

Next, the exogenous application of cytokinin in low nanomolar concentrations negatively affects the primary root growth and lateral root (LR) development. The loss-of-function mutant *cre1-2* has substantially decreased cytokinin sensitivity as demonstrated by normal primary root growth and lateral root development upon cytokinin application (Inoue et al., 2001). The primary root length and LR initiation thus became typical phenotypes followed in response to cytokinin. The primary root of CRE1-GFP lines was drastically shortened compared to *cre1-2* mutant and GFP/*cre1-2* control plants when grown on a medium supplemented with 20 nM BAP (Sup. Fig. 2A, B). Similarly, the CRE1-GFP lines and Col-0 developed no lateral roots when grown on cytokinin-containing medium. These results prove that the *pCRE1::CRE1-GFP* construct can fully complement the *cre1-2* mutant phenotype (Sup. Fig. 2A). On the other hand, the CRE1^int7^-GFP expressing seedlings (in Col-0 background) were less cytokinin sensitive when grown on medium with 20 nM BAP (Fig. 4A). Likewise, CRE1^int7^-GFP produced LRs analogous to *cre1-2* (Fig. 4C). CRE1^int7^-GFP plants showed increased primary root growth on the mock media during the early phases of seedling growth when compared to the Col-0, GFP/Col-0 control lines, and *cre1-2* (Sup. Fig. 3A, B). Moreover, the expression of two cytokinin primary target genes, *ARR5* and *CKX3*, was significantly lowered in the CRE1^int7^-GFP line compared to Col-0 (Sup. Fig. 4), further supporting the role of CRE1^int7^ in CKs signaling attenuation.

**Fig. 4.**
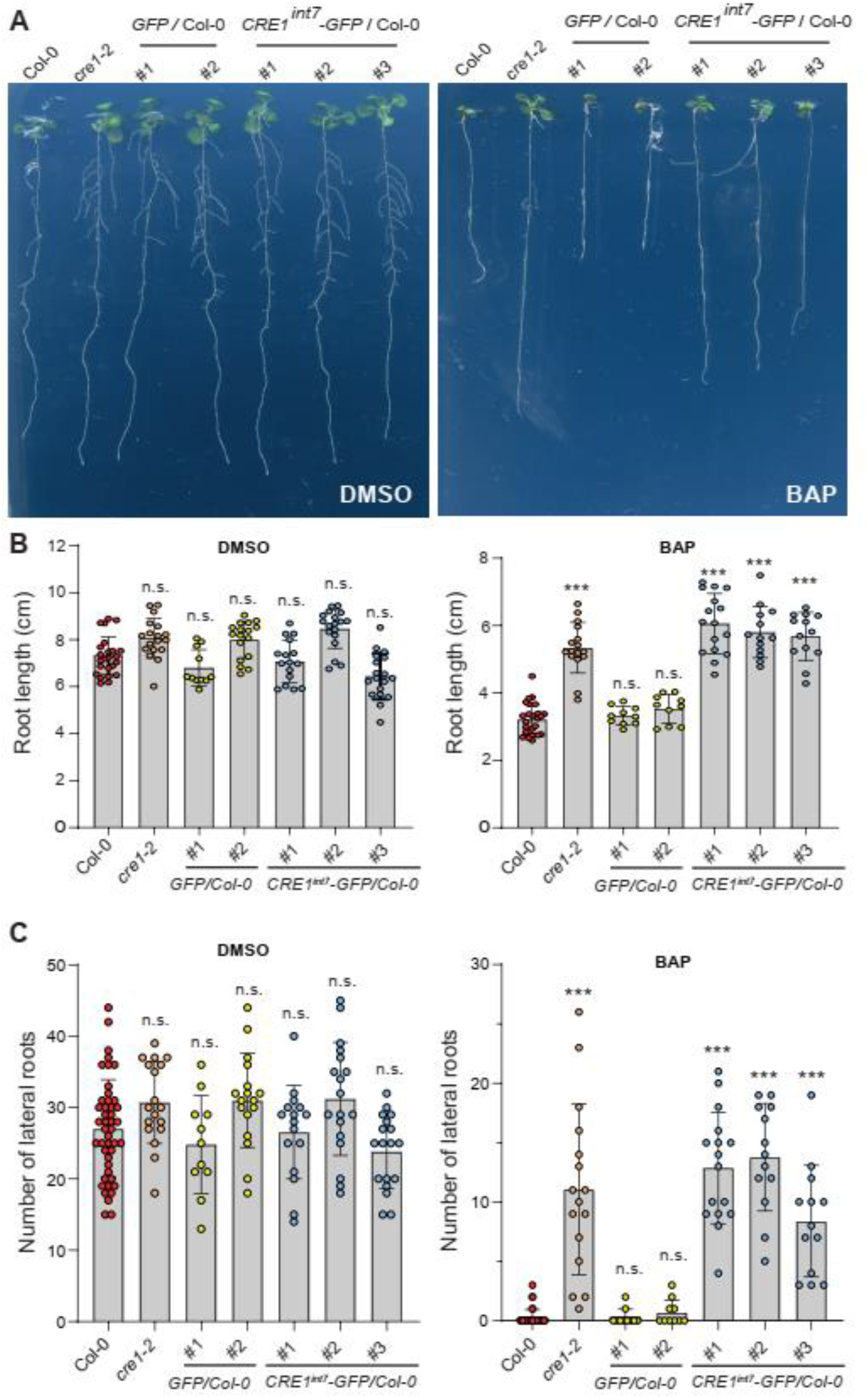
*Arabidopsis* lines expressing CRE1^int7^-GFP are less sensitive to cytokinin. The phenotype of seedlings expressing GFP alone (two independent lines) or CRE1^int7^-GFP (three independent lines) in Col-0 background. Seedlings were grown for 3 days on MS media and then transferred on media supplemented with cytokinin (20 nM BAP) or DMSO (mock control) and grown for another 14 days. **A**, Seedlings transferred on MS supplemented with cytokinin or DMSO. **B**, Primary root length was measured 14 days after transfer onto BAP/DMSO-containing media (17 days old seedlings). **C**, The number of emerged lateral roots was counted 14 days after transfer to BAP/DMSO-containing media (17 days old seedlings). Wild-type (Col-0) and mutant (*cre1-2*) are positive and negative controls for cytokinin treatment. Results represent means ± standard deviation (n ≥ 10). To compare the differences between groups the non-parametric Kruskal-Wallis ANOVA followed by multiple comparisons of mean ranks was used. Data comparing differences between Col-0 wild-type and individual mutants are shown (n.s. non-significant, *** p < 0.001).

Transgenes often undergo epigenetic silencing through mechanisms of Transcriptional Gene Silencing (TGS) or Post-Transcriptional Gene Silencing (PTGS). Both mechanisms can lead to *cis*-inactivation of transgene or *trans*-inactivation of the endogenous homologous genes reviewed in (Rajeevkumar et al., 2015). To investigate whether the phenotype of CRE1^int7^-GFP plants is not caused by the silencing of the endogenous *CRE1* gene, we monitored the expression of the native *CRE1* transcripts in the CRE1^int7^-GFP line and found it to be unchanged (Sup. Fig. 4). This shows that observed phenotype of CRE1^int7^ expressing line is not affected by transgene silencing.

Taking together, canonical CRE1-GFP and decoy CRE1^int7^-GFP receptors are co-expressed in the same tissue and subcellular compartments in *Arabidopsi*s. Moreover, the CRE1^int7^-GFP line show reduced cytokinin sensitivity, supporting the CRE1^int7^ decoy receptors’ function to attenuate the cytokinin signaling pathway.

## Discussion

The concept of a decoy receptor is a common strategy to regulate the sensing of primary pro-inflammatory cytokines and chemokines in mammals (McMahan et al., 1991). The canonical cytokine receptors are anchored in the membrane through a hydrophobic domain. After its proteolytic- or phospholipase C-mediated cleavage, truncated decoy receptors become soluble (Mantovani et al., 2001). Soluble receptors are released from the membrane, secreted outside the cell, and ready to titrate their ligand. This mechanism prevents the ligand from binding to the canonical cytokine receptors, so the cascade cannot be activated. The removal of the anchoring domain can be accomplished via alternative splicing as well (Vorlová et al., 2011).

Similarly, we discovered that AS acts on the *Arabidopsis* cytokinin receptor CRE1 and creates a novel functional *CRE1^int7^* variant. It retains the seventh intron within the transcript, introducing a premature termination codon thus encoding a truncated receptor (Fig. 1A). Despite the tendency of intron-retaining mRNA to be degraded via NMD (de Lima Morais and Harrison, 2010), it is becoming more clear that AS increases protein diversity (Wright et al., 2022) and is engaged in vivid metabolisms (Lam et al., 2022). We found that *CRE1^int7^* accumulates upon cytokinin application and decreases in cytokinin biosynthesis mutants (Fig. 1B, 1C). This transcript is not targeted to the NMD pathway (Fig. 1D) and its responsiveness to cytokinin (Fig. 1C) could be assumed as a part of the feedback regulation pathway. Our findings strongly suggest that CRE1^int7^ can modulate the cytokinin signaling by competing with the canonical CRE1 for ligand binding. This is the common mechanism of action by which decoy receptors work in mammalian cells (Mantovani et al., 2001).

Although the decoy receptor phenomenon has not yet been described *in planta*, we can find some analogical fine-tuning mechanisms in plant hormone pathways acting downstream of the receptors. These mechanisms involve the IR transcripts encoding truncated proteins often counteracting the function of the full-length proteins to modulate the signal pathway. Such a mechanism was recently described for jasmonate and abscisic acid signaling. Transcription factors of jasmonic acid signaling, *JAZ* genes, undergo IR producing truncated proteins that attenuate this pathway (Chung et al., 2010). A similar mechanism can be found in the ABA signaling pathway. The IR in the *HAB1* gene, encoding a regulatory phosphatase, generates a new form of protein with the opposite effect to full-length protein (Wang et al., 2015). The alternative splicing of the *HAB1* gene represents a mechanism to turn the ABA signaling on and off according to external cues (Wang et al., 2015).

Besides the established negative feedback mechanisms involving cytokinin-inducible (i) CKX expression controlling the ligand level, (ii) negative regulators AHP6 controlling signal transduction, and (iii) A-ARRs controlling gene transcription, the proposed CRE1^int7^ decoy receptors’ action represents another layer of fine-tuning control of the cytokinin signaling coming from cell compartmentalization.

The subcellular localization of the cytokinin receptors is a continuous dispute between exclusively PM (Inoue et al., 2001; Kim et al., 2006) and ER, with only a minor fraction residing at the PM (Caesar et al., 2011; Wulfetange et al., 2011). Even though only a small portion of the cytokinin receptor pool localizes at the PM, it appears to be of importance. The depletion of the cytokinin pool in the apoplast has a major effect on cytokinin perception, while depletion of the cytosolic pool has not (Zürcher et al., 2016). Recently, Antoniadi and colleagues came to a similar conclusion using immobilized CK ligands which activate cytokinin signaling although the ligands cannot pass through the PM (Antoniadi et al., 2020).

Our novel *pCRE1::CRE1-GFP* line, which is the first CRE1 reporter line complementing the *cre1* mutant phenotype, as far as we are concerned, as well as the CRE1^int7^-GFP-expressing line, have repeatedly localized both CRE1 proteins to PM and ER of root stellar cells (Fig. 3A). It corresponds with previously reported CRE1 localization studied in root epidermal cells driven by *CaMV35S* promoter (Kubiasová et al., 2020). CRE1 likely undergoes subcellular trafficking, and the receptor distribution is not static (Kubiasová et al., 2020). The broad subcellular localization suggests that CRE1 might either function in multiple subcellular domains or rather be compartmentalized to modulate its activity. In the context of recently published works showing the PM being the dominant site of CK perception (Zürcher et al., 2016; Antoniadi et al., 2020), we assume the second scenario seems to be likely. Consequently, both CRE1 variants, occupying the same compartments, compete for ligands to either activate or attenuate the cascade. Due to CRE1 dual localization, we cannot exclude the possibility that cytokinin perception occurs in ER to some extent as well.

### Conclusions and Perspectives

The principle of DcRs counteracting the activity of canonical receptors is becoming a legitimate therapeutical strategy to treat virus infections and genetically encoded diseases including cancer and inflammatory diseases (Gershoni, 2008). The engineered decoy receptors were successfully used to neutralize the virus particles (Capon et al., 1989; Arimori et al., 2022) or apoptosis-inducing factors (Kariolis et al., 2014; Miao et al., 2022).

Cytokinin is one of the crucial morphogens of root architecture controlling root meristem activity. An enhanced root system is one of the key traits for breeding drought stress-tolerant crops. Nevertheless, the agricultural use of cytokinins has many limitations including strong inhibitory effects on primary root growth and lateral root initiation (Laplaze et al., 2007). Genetically engineered CRE1^int7^ decoy receptors could serve as an elegant tool to fine-tune the cytokinin signaling in a tissue-specific and/or spatiotemporal manner to achieve interesting root phenotypes with agricultural implications. We believe the concept of decoy receptors could be extrapolated to other plant hormone signaling pathways as well.

## METHODS

### Plant materials and growth conditions

All mutants used in the study were Col-0 ecotypes that served as wild-type control. *cre1-2*, *ipt2/9*, and *ipt3/5/7* mutants were provided by Dr. Tatsuo Kakimoto (Osaka University, Japan). *upf1-5* and *upf3-1* were provided by Dr. Nicola Cavallari (ISTA, Austria).

The seeds were surface-sterilized and sown on standard Murashige and Skoog (MS) medium (0.43% MS, 1% sucrose, 1% plant agar, from Duchefa, cat. no. P1001) plates and stratified at 4 °C in dark for 2-3 days. Seedlings were then grown on vertically oriented plates in growth chamber at 21°C under long-day conditions (16 h light of 120 μmol m^-2^ s^-1^/8 h dark), if not stated otherwise.

### Cloning and construct preparation

The sequences of all primers used in this work are listed in the Sup. Tab. 1. For bacterial assays in *E. coli*, the original *pIN-III* expression vector was first modified by inserting a *4xMyc* tag sequence, followed by a *tNOS* terminator. The resulting construct was named *pIN-III_4xMyc_tNOS*. Open reading frames (ORFs) of *CRE1* (primers: CRE1_ATG1_Fw_BamHI and CRE1_WT_rev_SpeI) and *CRE1^int7^* (primers: CRE1_ATG1_ Fw_BamHI and CRE1int7_rev_SpeI) were PCR amplified from *Arabidopsis thaliana* Col-0 root cDNA and cloned into *pIN-III 4xMyc_tNOS* under BamHI and SpeI restriction sites.

To generate constructs for transient expression in protoplasts and TCS::LUCIFERASE (TCS::LUC) assay, the *eGFP* reporter gene was subcloned (primers: eGFP_Fw_EagI and GFP_rev_EagI_XhoI_EcoRV_XbaI) into the *pENTR2B* vector using EagI restriction enzyme. The resulting construct *pENTR2B_GFP* has a unique NotI site located upstream of *eGFP* ORF. The *CRE1* (primers: CRE1b_fw_SalI and CRE1abc_rev_NotI) and *CRE1^int7^* (primers: CRE1b_fw_SalI and CRE1bint7_R_NotI) ORFs were PCR amplified and cloned into *pENTR2B_GFP* under SalI and NotI restriction sites to allow for C-terminal fusion with *eGFP*. The *CRE1-GFP* fusions were shuttled into the *p2GW7,0* expression vector using Gateway LR reaction to generate *35S::CRE1-GF*P constructs.

The ER marker construct *35S::AtWAK2-tdTomato-HDEL* was provided by Dirk Becker (University of Hamburg, Germany).

The *pENTR2B::CRE1/AHK4-eGFP* construct containing a genomic fragment of *CRE1* fused with an *eGFP* reporter (Kubiasová et al., 2020) was used to generate the *pCRE1::CRE1-GFP*. The *TMVΩ* translational enhancer sequence was introduced just upstream from the first ATG using the oligo annealing method (TMV_oligo_F_SalI; TMV_oligo_R_SalI) into the SalI restriction site. Resulting construct *pENTR2B::TMVΩ_gCRE1-eGFP* served as a template for preparing the *pENTR2B::TMVΩ_gCRE1^int7^-eGFP* construct by inversion PCR method using phosphorylated primers. The primers gCRE1_int7_Rv and Linker_fw were used. The PCR product was digested by DpnI enzyme to remove the template plasmid, purified using NucleoSpin Gel and PCR Clean-up kit (Macherey-Nagel), and circularized using T4 DNA ligase.

Since the original GFP-only control construct did not show any fluorescence *in planta*, the first *CRE1* exon and intron sequences and the beginning of the second exon were included. The control GFP-only construct was generated by inversion PCR on *pENTR2B::TMVΩ_gCRE1-eGFP* (primers: GFP_GA2_FW; pEN_GFP_CTRL_GA_RV) to skip the region between second and last exons of *CRE1* gene. The PCR product was digested by the DpnI enzyme to remove the template plasmid, gel-purified, and self-assembled using Gibson assembly (NEBuilder^®^ HiFi DNA Assembly Master Mix, NEB).

All resulting *pENTR2B* constructs were verified by sequencing, then shuttled into the *pWOL::GW* vector (Michniewicz et al., 2015) carrying the 2 kbp promoter region of the *WOL/CRE1/AHK4* gene using Gateway LR reaction (ThermoFisher), and sequenced.

Sequences and maps of all plasmid constructs prepared in this work are provided as SnapGene files in the Supplement.

### Plant transformation

Transgenic *Arabidopsis* plants were generated by the floral dip method using *Agrobacterium tumefaciens* strain GV3101 (Clough and Bent, 1998). Transformed T1 seedlings were selected in soil by spraying with 0.02% BASTA solution. The following transgenic generations were selected on a medium supplemented with 15 μg mL^-1^ phosphinothricin.

### Quantitative PCR with reverse transcription

Total RNA was extracted from roots of 4-day-old untreated seedlings (Fig. 1C, D; Sup. Fig. 4) and seedlings sprayed with mock (DMSO) or 1 μM BAP for 60 min (Fig. 1B) using RNAqueous™ Total RNA Isolation Kit (ThermoFisher).

RNA was treated twice with a TURBO DNase-free™ kit (ThermoFisher). The presence of genomic DNA contamination was tested with the specific primer pair annealing in the promoter region of the *ERF8* gene (Ethylene Response Factor 8; primers ERF8_TPF; ERF8_TPR).

Poly(dT) cDNA was prepared from 3 μg total RNA with RevertAid H Minus Reverse Transcriptase (ThermoFisher) and analyzed on StepOnePlus Real-Time PCR System (LifeTechnologies) with gb SG PCR Master Mix (Generi Biotech) according to the manufacturer’s instructions. The expression of all *CRE1* transcripts was quantified using a primer pair specific to the last *CRE1* exon (CRE1b_exon10_fw and CRE1b_exon10_rev). The *CRE1^int7^* transcript was quantified using a primer pair annealing to exon7 and intron 7 (CRE1b_exon7_fw and CRE1b_int7_rev). The *ARR5* and *CKX3* expression was quantified using primer pairs (ARR5_fw and ARR5_rev; RT-CKX3_F and RT-CKX3_R). The relative expression of respective targets was normalized to the housekeeping gene *TUBULIN 3* (TUB3_F2426 and TUB3_R2616). The splicing index of the seventh *CRE1* intron was calculated as the *CRE1^int7^* expression normalized to all *CRE1* transcripts (primers CRE1b_exon10_fw and CRE1b_exon10_rev). The primer position within the *CRE1* transcript is shown in Fig. 1A. For cytokinin treatment experiments, the fold change refers to mock treatment. In the case of the mutant experiment with *ipt* and *upf* mutants, as well as with the *pCRE1::CRE1^int7^-GFP* line, the fold change refers to Col-0 wild-type.

Plant material for cytokinin treatment (Fig. 1B) was prepared and harvested on separate days in a total of nine independent experiments. RT-qPCR data were collected from five technical replicates. In the case of *ipt, upf* mutants (Fig. 1C, D), and *pCRE1::CRE1^int7^-GFP* line (Sup. Fig. 4), five biological replicates were used. RT-qPCR was performed in three technical replicates.

### Microscopy

Zeiss LSM800 and Zeiss LSM900 confocal scanning microscopes equipped with 40x Plan-Apochromat water immersion objective were employed to follow the expression of the fluorescent reporters. Images were captured using excitation wavelengths of 488 nm for eGFP; 499 nm for Alexa Fluor 488; 555 nm for Cy3; 561 nm for tdTomato and propidium iodide. Zeiss Zen Blue and ImageJ software version 1.53c were used for image postprocessing.

### Ligand-binding assay in *E. coli*

The receptor binding assay was performed using the *E. coli* strain KMI001 clones carrying the *pIN-III* constructs expressing 4xMyc-tagged *CRE1* or *CRE1^int7^*. The expression of CRE1 proteins was induced with 0.25 mM IPTG in cultures of OD600 (0.6 – 0.7) and then cultivated for 5h at 18 °C. The competitive binding assay was performed as described in (Kubiasová et al., 2020). The competition reaction was allowed to proceed with 3 nM [2-^3^H]*t*Z and 10 μM of cold iP and *t*Z, or 0.1% (v/v) DMSO (mock control). [2-^3^H]*t*Z was provided by Dr. Zahajská from the Isotope Laboratory, Institute of Experimental Botany, Czech Academy of Sciences.

### Receptor activation assay in *E. coli*

The receptor activation assay was performed as described previously (Suzuki et al., 2001). The same *pIN-III* clones of *E. coli* strain KMI001 used in the ligand-binding assay were also used for this assay. The expression of CRE1 proteins was induced with 0.25 mM IPTG in cultures of OD600 (0.6 – 0.7) and then cultivated for 16 h at 18 °C. The induced culture was diluted to OD600 (0.2) and the experiment was performed in 96-well plates for 6 h at 18 °C in M9 media containing 1 μM iP or DMSO (mock).

### Transient expression in protoplasts and TCS::LUC assay

The transient expression was performed in protoplasts isolated from 4-day-old *Arabidopsis* Col-0 root suspension culture. Protoplasts were transfected by PEG as described previously (Hurný et al., 2020). The quality of protoplasts was checked under the microscope, and density was quantified by spectrophotometry (OD600). Protoplasts were diluted in B5 glucose-mannitol solution to 1.6 OD600. 50 μL of protoplasts were co-transfected with a 2 μg ER marker (*35S::AtWAK2-tdTomato-HDEL*) and 5μg of *p2GW7.0* plasmid DNA (*35S::CRE1-GFP, 35S::CRE1^int7^-GFP* or *35S:: GFP*). The transient expression of CRE1-GFP proteins was allowed for 16 h after PEG transfection in the dark at RT. Transfected protoplasts were mounted on the Nunc™ Glass Bottom Dishes (Thermo Fisher, cat. no. 150680) and observed with Zeiss LSM 800 laser scanning confocal microscope.

The expression of CRE1-GFP proteins was verified by Western blotting with an anti-GFP antibody on microsomal fractions isolated from protoplasts according to the protocol described in (Hajný et al., 2020). In the case of protoplast expressing the eGFP alone, the protein was precipitated from soluble protein fraction by methanol/chloroform procedure. Briefly, a mixture of 200 μL of chloroform, 800 μL methanol, and 600 μL of Milli Q water was added to 200 μL of soluble protein fraction, vortexed, and centrifuged at 13 000 rpm at RT for1 min. The upper phase was discarded and 600 μL of fresh methanol was added to the mixture, vortexed, and centrifuged for 3 min. The supernatant was discarded. Dried pellet was resuspended in 50 μL of protein extraction buffer (PEB, 50 mM Tris, 150 mM NaCl, 0.5 mM EDTA, PhosSTOP, cOmplete™, EDTA-free Protease Inhibitor Cocktail) with detergents (0.5% Triton X-100; 0.5% CHAPS) and analyzed on SDS-PAGE.

The luciferase transient expression assay (TCS::LUC) was performed on protoplasts isolated from 4-days-old *Arabidopsis* Col-0 root suspension culture as described previously (Kubiasová et al., 2020). Protoplasts were cotransfected with 2.5 μg of TCS::LUC a cytokinin reporter plasmid (Müller and Sheen, 2008) expressing Firefly luciferase (*f*LUC), 2.5 μg of normalization plasmid containing the *Renilla* luciferase (*r*LUC), and 5 μg of *p2GW7,0* plasmid carrying either cytokinin receptor (*35S::CRE1-GFP* and *35S::CRE1^int7^-GFP*) or GFP only (*35S::GFP*) constructs.

### Western blotting

For immunoblot analysis, 20 μL of microsomal protein fractions (*35S::CRE1-GFP* and *35S::CRE1^int7^-GFP*) or soluble protein fractions (for *35S::GFP*) isolated from transfected protoplasts were separated by SDS-PAGE and subjected to immunoblotting using an anti-GFP-HRP antibody (130-091-833, Miltenyi Biotec, dilution 1:5000).

In the case of expression in *E.coli*, the bacterial pellet was resuspended in 1x Laemmli buffer, boiled at 95 °C for 5 min, and centrifuged at 14 000 rpm. 5 μL of supernatant was analyzed by SDS-PAGE and subjected to immunoblot analysis with mouse anti-Myc (clone 4a6, Millipore, cat. no. 05-724, dilution 1:1000) and goat anti-mouse IgG-HRP (Santa Cruz Biotechnology, cat.no. sc-2005, dilution 1:5000).

### Root phenotypes

Root growth and lateral root number were assayed using seedlings grown vertically on a 1xMS medium. Seedlings were grown on MS media for three days, then transferred to new MS plates supplemented with 20 nM BAP or DMSO (mock control) and grown for another 10 days. Images were taken with an EPSON perfection v800 Photo scanner at selected time points. The root length was evaluated using 5-day-old seedlings (n ≥ 16). Lateral root density quantification was performed using 14-day-old seedlings (n ≥ 16). The statistical significance was evaluated with the Student’s t-test and two-way ANOVA.

### Statistical analysis

The statistical significance was evaluated with the Student’s t-test and two-way ANOVA.

## Acknowledgments

We dedicate this paper to the deceased P. Galuszka for his inspiration and support of our project. This research was supported by the Scientific Service Units (SSU) of IST-Austria through resources provided by the Imaging and Optics Facility (IOF), and the Life Science Facility (LSF). The work was supported by Czech Science Foundation via project 18-23972Y (D.Z., I.K., K.K.).

## Author contributions

Y.I., E.B., and D.Z. initiated the project and designed the experiments. I.K., Y.I., and D.Z. conducted RT-qPCR and 5’RACE. J.H. and D.Z. performed microscopic experiments. K.K. made ligand-binding and receptor activation assay. M.G. edited figures. O.N. analyzed the data. H.S. and D.Z. made protoplast experiments. M.K. performed most of the *Arabidopsis* work, phenotyping, and statistical analysis. I.K., J.H., and D.Z. wrote the manuscript. All authors read and edited the manuscript.

---

**Sup. Fig. 1.**
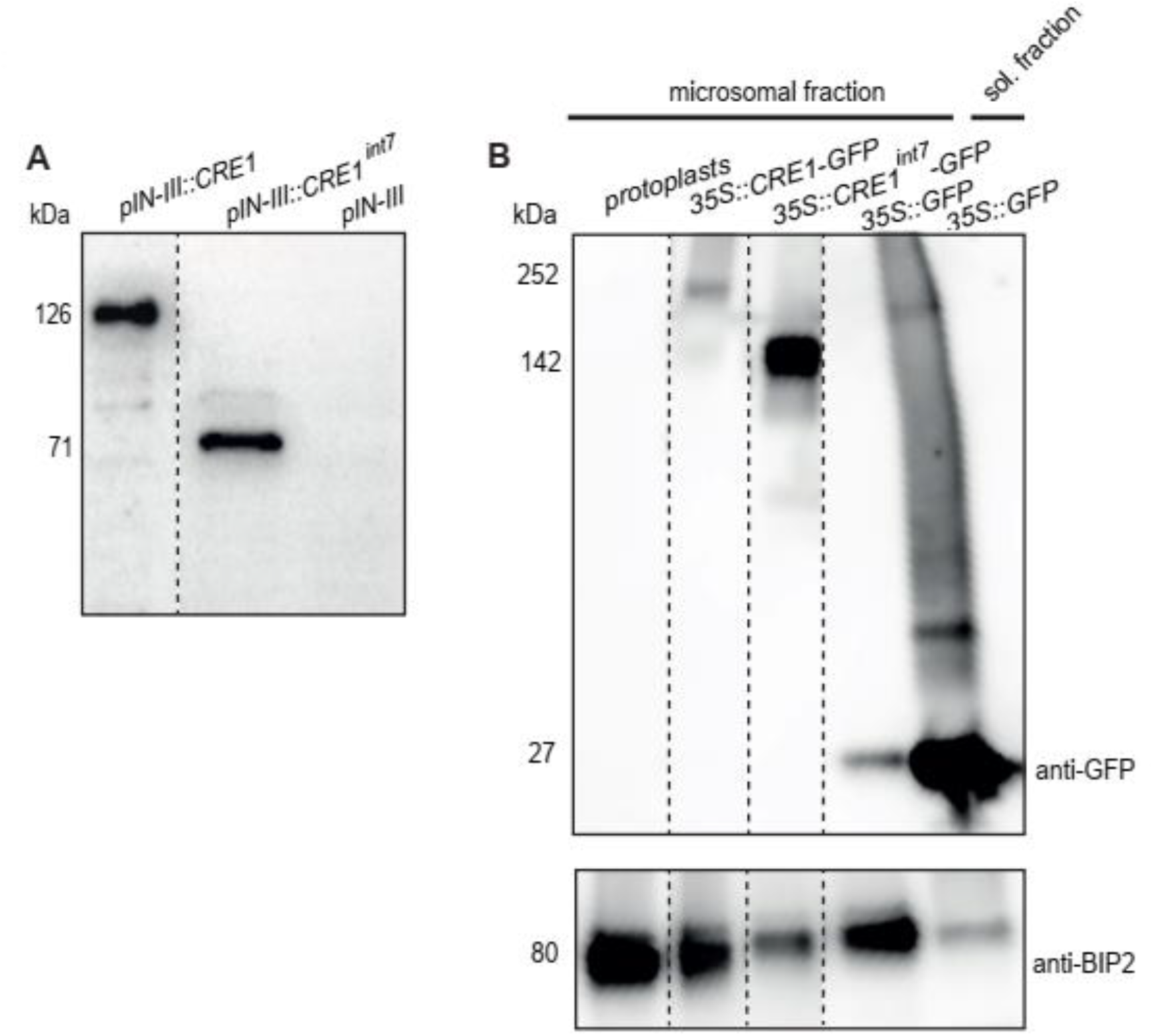
CRE1 and CRE1^int7^ expressed in *E.coli* and *Arabidopsis* protoplasts form stable proteins. **A,** Western blot analysis of total protein extract from *Escherichia coli* expressing CRE1-4xMyc and CRE1^int7^-4xMyc. Clone carrying empty plasmid *pIN-III* was used as control. These clones were used for both ligand-binding and receptor activation assays. Membranes were incubated with an anti-Myc antibody. The molecular weight of CRE1-4xMyc (126 kDa) and CRE1^int7^-4xMyc (71 kDa) corresponds to the expected molecular weight of receptor monomers. **B,** Western blot analysis of microsomal fractions of protoplasts expressing CRE1-GFP and CRE1^int7^-GFP. Protoplasts expressing GFP alone (*35S::GFP*) and non-transformed protoplasts (protoplasts) were controls. Since GFP is soluble, the soluble protein fraction was included for *35S::GFP*. Membranes were incubated with anti-GFP and anti-BIP2 (loading control) antibodies. The molecular weight of CRE1-GFP (252 kDa) and CRE1^int7^-GFP (142 kDa) corresponds to the predicted molecular weight of receptor dimers. The molecular weight of GFP alone and BIP2 is 27 kDa and 80 kDa, respectively. The non-adjacent lanes were re-arranged and the boundary between lanes is delineated by a dashed line.

**Sup. Fig. 2.**
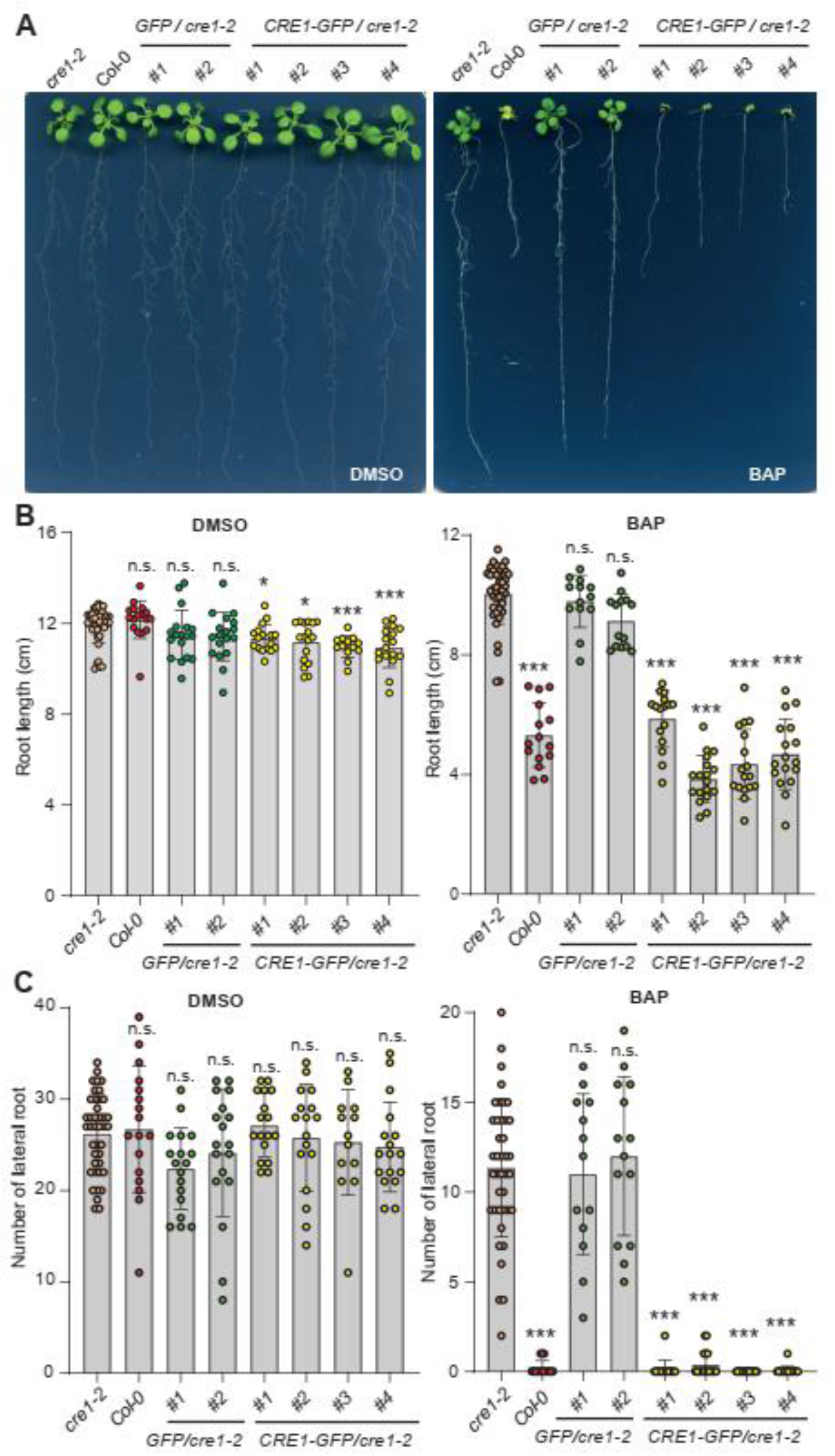
Novel *pCRE1::CRE1-GFP* construct rescues *cre1-2* mutant phenotype. The phenotype of seedlings expressing GFP alone (two independent lines) or CRE1-GFP (four independent lines) in the *cre1-2* mutant background in response to cytokinin. Seedlings were grown for 3 days on MS media, then transferred on media supplemented with cytokinin (20 nM BAP) or DMSO (mock control) and grown for another 11 days. **A**, Seedlings transferred on MS supplemented with cytokinin or DMSO. **B**, Primary root length was measured 9 days after transfer onto BAP/DMSO-containing media. **C**, The number of emerged lateral roots was counted 11 days after transfer to BAP/DMSO-containing media (14 days old seedlings). Wild-type (Col-0) and mutant background (*cre1-2*) are positive and negative controls for cytokinin treatment. Results represent means ± standard deviation (n ≥ 10). To compare the differences between groups the non-parametric Kruskal-Wallis ANOVA followed by multiple comparisons of mean ranks was used. Data comparing differences between *cre1-2* and individual transgenic lines are shown (n.s. non-significant, * *P* < 0.05, *** p < 0.001).

**Sup. Fig. 3.**
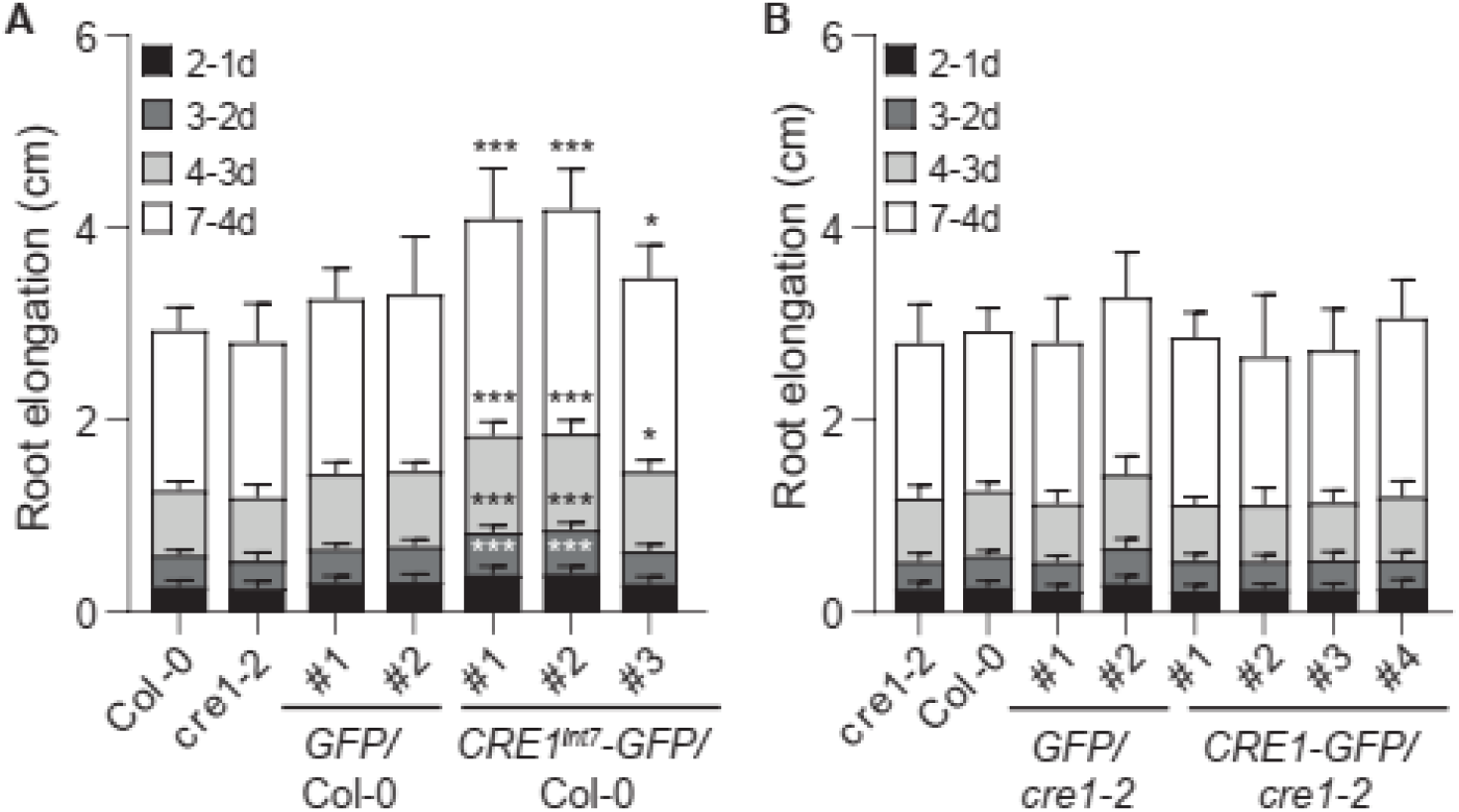
*Arabidopsis* lines expressing CRE1^int7^ show accelerated root growth. Primary root growth of *Arabidopsis* lines expressing CRE1^int7^-GFP (**A**) and CRE1-GFP (**B**) was measured at selected time points. Seedlings were grown vertically on MS media without cytokinin. Col-0, *cre1-2, GFP*/Col-0 and *GFP/cre1-2* are controls. Three independent lines (*CRE1^int7^-GFP*), four independent lines (*CRE1-GFP*), and two independent lines (*GFP*/Col-0 and *GFP/cre1-2*) were analyzed. Results represent means ± standard deviation (n ≥ 20). To compare the differences between groups the non-parametric Kruskal-Wallis ANOVA followed by multiple comparisons of mean ranks was used. Data comparing differences in delta root length (2-1d, 3-2d, 4-3d, and 7-4d) between Col-0 wild-type and individual transgenic lines are shown (* *P* < 0.05, *** *P* < 0.001).

**Sup. Fig. 4.**
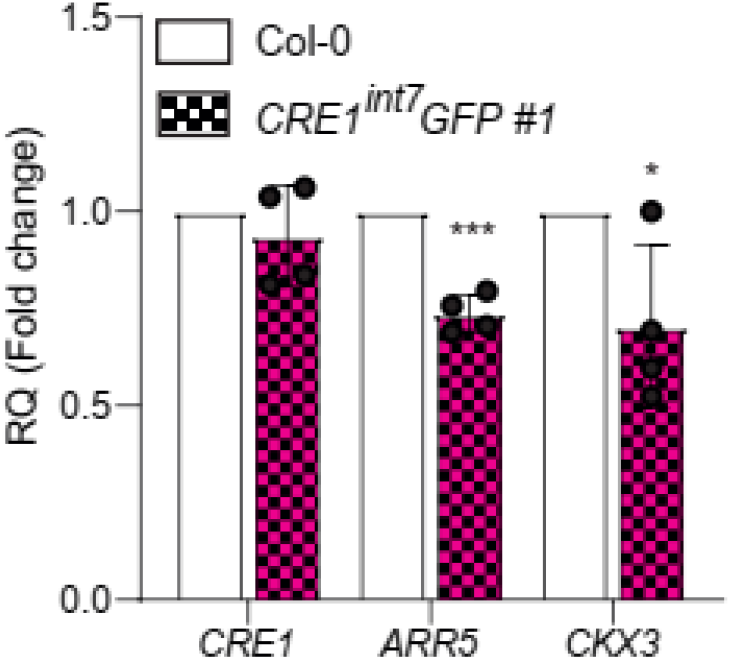
Cytokinin signaling is attenuated in *Arabidopsis* lines expressing CRE1^int7^, while endogenous *CRE1* expression is unaffected. Quantitative RT-PCR of endogenous *CRE1* and two cytokinin primary responsive genes *ARR5* and *CKX3* in CRE1^int7^-GFP-expressing line. The relative gene expression (RQ) was normalized to *TUBULIN3* in wild-type (Col-0). Values show the mean from four biological repeats ± standard error. The *P*-value shows the Student’s t-test, data comparing differences between Col-0 wild-type and CRE1^int7^-GFP-expressing line. \**P* < 0.05, \*\*\**P* < 0.001.

**Sup. Tab. 1.**
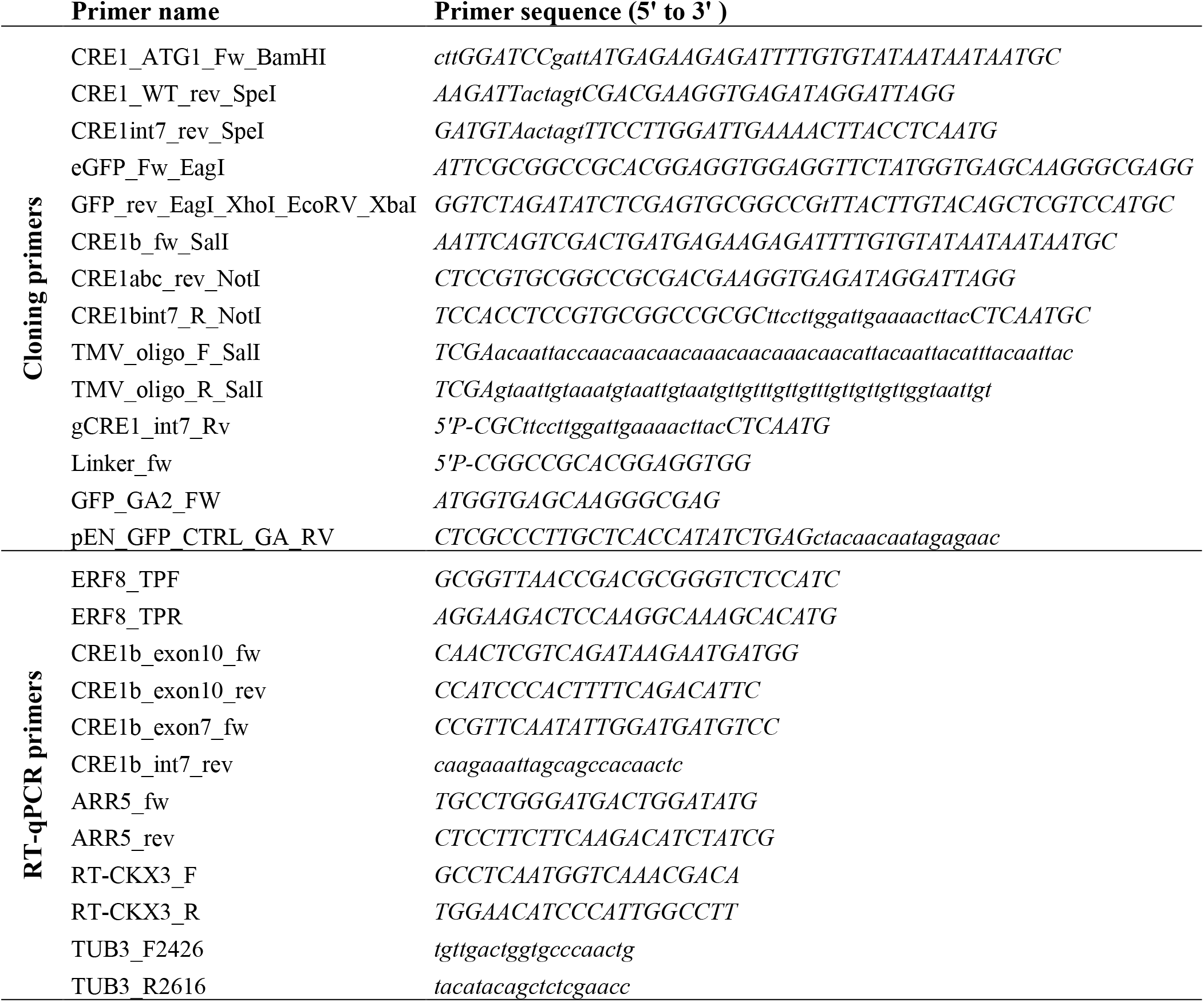
Primer list.

## Notes

### Competing Interest Statement

The authors have declared no competing interest.

